# Scans of the *MYC* mRNA reveal multiple stable secondary structures—including a 3′ UTR motif, conserved across vertebrates, that can affect gene expression

**DOI:** 10.1101/564864

**Authors:** Collin A. O’Leary, Ryan J. Andrews, Van S. Tompkins, Jonathan L. Chen, Jessica L. Childs-Disney, Matthew D. Disney, Walter N. Moss

## Abstract

The *MYC* gene encodes a human transcription factor and proto-oncogene that is dysregulated in over half of all known cancers. To better understand potential post-transcriptional regulatory features affecting *MYC* expression, we analyzed secondary structure in the *MYC* mRNA using a program that is optimized for finding small locally-folded motifs with a high propensity for function. This was accomplished by calculating folding metrics across the *MYC* sequence using a sliding analysis window and generating unique consensus base pairing models weighted by their lower-than-random predicted folding energy. A series of 30 motifs were identified, primarily in the 5’ and 3’ untranslated regions, which show evidence of structural conservation and compensating mutations across vertebrate *MYC* homologs. This analysis was able to recapitulate known elements found within an internal ribosomal entry site, as well as discover a novel element in the 3’ UTR that is unusually stable and conserved. This novel motif was shown to affect *MYC* expression: likely via modulation of miRNA target accessibility. In addition to providing basic insights into mechanisms that regulate *MYC* expression, this study provides numerous, potentially druggable RNA targets for the *MYC* gene, which is considered “undruggable” at the protein level.

## Introduction

The *MYC* proto-oncogene is an important transcription factor that is required for programmed cell death (apoptosis) and cell proliferation [1]. It is a key component of oncogenesis [2] and, indeed, *MYC* is dysregulated in >50% of all cancers [3]. Post-transcriptional control plays significant roles in the regulation of many genes includin*g M*YC. Within the 5’ <u>u</u>ntranslated region (UTR) of the *MYC* mRNA is a structured internal ribosomal <u>e</u>ntry <u>s</u>ite (IRES) that stimulates cap-independent translation under conditions where cap-dependent translation is inhibited: e.g. during apoptosis [4]. Consistent with other IRESs [5] the *MYC* IRES secondary structure, deduced from *in vitro* chemical probing data [6], is complex and contains two pseudoknots—motifs comprised of “non-nested” base pairing between looped out regions of RNA [7]. In addition to the IRES, other post-transcriptional regulatory mechanisms affect MYC expression—e.g. <u>mi</u>cro<u>R</u>NAs (miRs) [8]—that may be affected by RNA structure [9].

To determine if other structured RNA regulatory elements can be playing roles in *MYC* expression, we applied a methodological pipeline for RNA motif discovery that was optimized from studies of the *Xist* lncRNA [10], as well as the Human [11], Zika and HIV genomes [12]. There are two major steps in this pipeline: (1) a scanning step, where the RNA is examined using a sliding analysis window to record predicted metrics important for analyzing RNA secondary structure (e.g. the thermodynamic stability); and, (2) an analysis step where unique local motifs are defined then evaluated vs. comparative sequence/structure and/or experimental probing data. Each step is achieved using the programs `ScanFold-Scan` and `ScanFold-Fold`, respectively. Used together these programs define the potential RNA structural properties of long sequences and identify motifs likely to be ordered to form, presumably functional, defined structures. This is accomplished by generating consensus structure models across all scanning windows, where base pairs are weighted by their thermodynamic z-score: a measure of the unusual stability of a sequence that is calculated by comparison to the folding energy of matched randomized control sequences. Here, negative values indicate sequences that are ordered to fold and that may be functional [13]. While the primary goal is to deduce what nucleotides may be functionally significant, `ScanFold-Fold` models can also increase prediction accuracy [12]

In this report, `ScanFold-Scan` and `ScanFold-Fold` were applied to the longest *MYC* RefSeq mRNA isoform to generate a map of its folding landscape as well as deduce motifs important to the regulation of expression. Numerous motifs were deduced, including those that recapitulated known structures in the *MYC* IRES.

## Results

### ScanFold-Scan mapping of secondary structure in the MYC mRNA

To predict RNA secondary structural characteristics important to *MYC* function, the RefSeq mRNA (NM_001354870) was analyzed using the program `ScanFold-Scan` [12]. This isoform was selected for analysis, as it would contain all potential structural elements found in other (shorter) *MYC* isoforms. The mRNA sequence was analyzed using a 1 nt step and 70 nt window size (Figure 1 and Document S1). Several folding metrics were calculated across analysis windows, which are described in detail in the Materials and Methods and in reference [12]. Briefly, the ΔG° measures the minimum (lowest or most stable) predicted change in the Gibb’s free energy upon RNA folding and indicates the thermodynamic stability of RNA structure. The <u>e</u>nsemble <u>d</u>iversity (ED) is a measure of the structural diversity predicted in the folding ensemble: low numbers indicate one or few dominant structures, while higher numbers indicate multiple conformations or a lack of structure. The z-score measures the propensity of a sequence to be ordered to fold into stable structures. Negative z-scores give the number of standard deviations more thermodynamically stable a sequence is vs. random (see Eq. 1).

**Figure 1.**
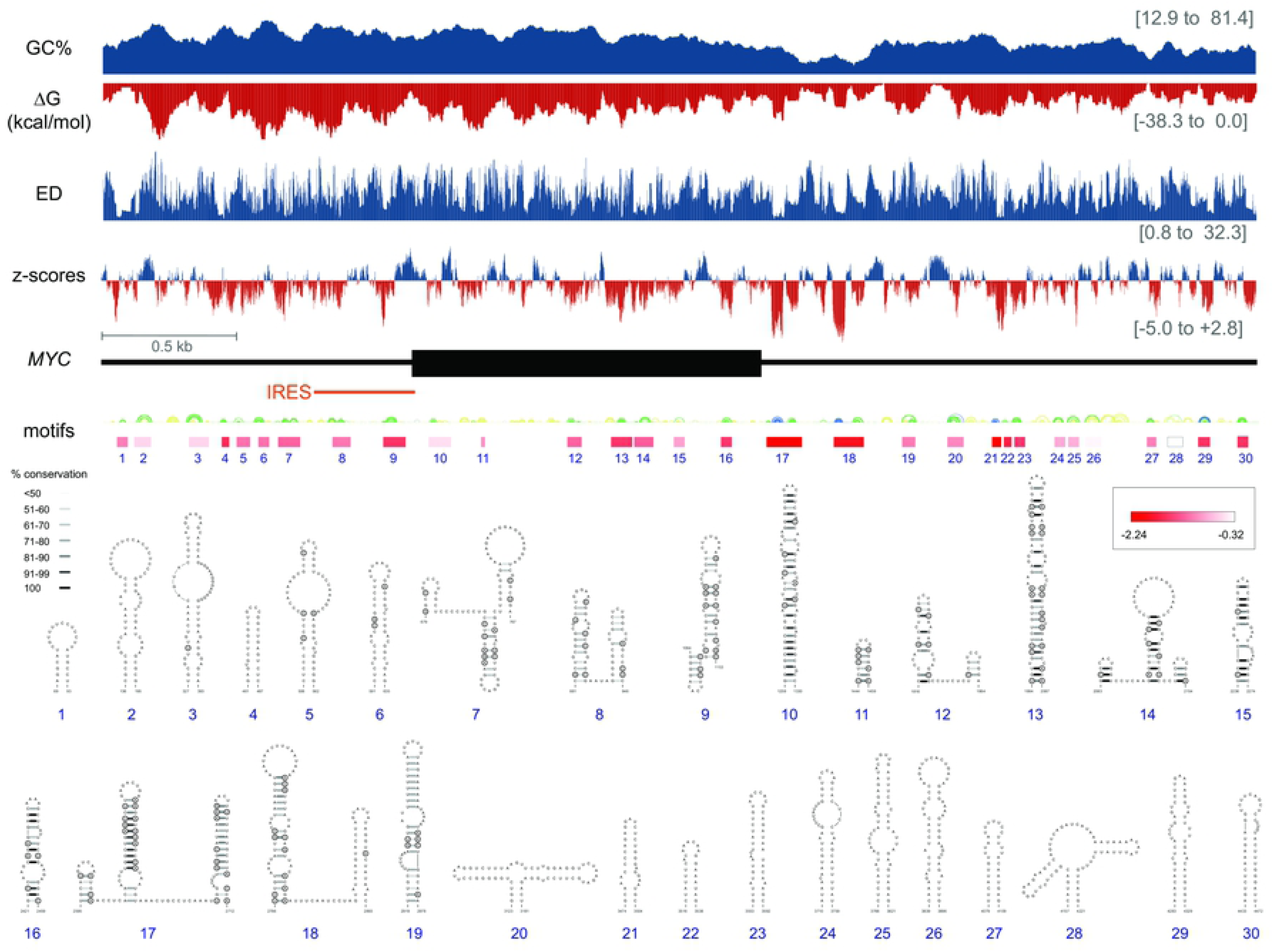
Summary of `ScanFold-Scan` and `ScanFold-Fold` results for the short UTR *MYC* mRNA. At the top are charts indicating the predicted `ScanFold-Scan` metrics across the mRNA. The bars are set at the 1^st^ nt of the 70 nt window, thus data corresponds to the 70 nt downstream of the bar. Below these is a cartoon of the *MYC* mRNA with UTRs and coding region represented in thin and thick black lines, respectively. This cartoon is annotated with boxes which depict the location and extent of `ScanFold-Fold` predicted motifs shaded red based on the average z-score of windows in which motif base pairs occurred. Below these are RNA secondary structure arc diagrams which depict the most favorable base pairs predicted via `ScanFold-Fold`, colored according the average z-scores of windows in which they appear (with blue, green and yellow corresponding to less than −2, −1 and 0 z-scores averages respectively). Below these, are refolded models of the motifs built with −1 average z-score bps as constraints. Each is annotated with their bp conservation as determined from an alignment of 15 representative mRNAs (Document S3) indicated by shading on the base pair (see key). Circled bases are sites of putative structure-preserving consistent and compensatory mutations.

The global trends in each metric are shown at the top of Figure 1. The trend in the predicted thermodynamic stability approximately follows the GC% and decreases across *MYC*: going from the highly stable 5’ UTR to the relatively unstable 3’ UTR. This trend is also discernible in the distributions of predicted ΔGs for windows spanning the 5’ UTR, coding region and 3’ UTR of the mRNA (Fig. S1). The windows spanning the 5’ and 3’ UTR junctions are less stable than flanking sequences. ED values are more evenly distributed across *MYC*, however, jumps in values for windows spanning the 5’ and 3’ UTR coding region junctions were observed (Figs. 1 and S1). Thermodynamic z-scores ranged from highly negative (−5.0; or 5 standard deviations more stable than random) values to positive ones (+2.8). The average z-score across *MYC* was only slightly negative (−0.4) and there was no evidence of global bias in z-score toward negative values. Notably, windows spanning the 5’ UTR junction were shifted toward *positive* z-scores (average of +1.4). These trends in RNA structural stability are more striking when considering “short” UTR isoforms for *MYC* (Fig. S2), which end just upstream of the *MYC* IRES and downstream of Motif 17 (Fig. 1).

### ScanFold-Fold prediction of functional RNA structural motifs

To deduce local RNA folding that may be functionally significant, all `ScanFold-Scan` prediction windows were analyzed using `ScanFold-Fold`. The `ScanFold-Fold` program generates weighted consensus secondary structures, where minimum free energy (MFE) base pairs that contribute to low z-scores are deduced across the scans. Using a cutoff of −1 `ScanFold-Fold` identified 354 bp (Document S2) across the mRNA, while a cutoff of −2 yields 46 bp that are localized to the 3’ UTR. Refolding the mRNA with −1 `ScanFold-Fold` bp as constraints added 153 bp to the discovered motifs: e.g. by extending helices or closing unpaired bases in the consensus prediction (Document S2). These 507 bp are divided into 30 motifs that span the *MYC* mRNA (Fig. 1). Motif locations, as expected, correspond to negative dips in z-score; however, dips in ΔG° and ED are also observed at motif sites. The most prominent regions with dips in metrics occur at Motifs 17 and 18 (Fig. 1), which contain very low z-score base pairs (cutoff < −2) deduced by `ScanFold-Fold`. These two motifs, particularly Motif 17, had the most favorable ScanFold metrics of any region/motif predicted for MYC.

All motif bp were analyzed versus an alignment of 15 vertebrate mRNA sequences (Document S3). Motif 17 had the highest conservation of structure and was supported by the greatest number of consistent and compensatory mutations (Fig. 1). In general, Motifs 7–19 showed evidence of conservation, however, little conservation data was found outside these regions: particularly downstream of Motif 19, where the long 3’ UTR annotated for the human *MYC* RefSeq mRNA is not present in the RefSeq mRNA annotations of other species (Document S3).

### Analysis of the MYC 5’ and 3’ UTRs

Motifs 8 and 9 overlap a previously-studied structural feature of the *MYC* mRNA: the IRES [4, 6]. Motif 8 is recapitulated in the *MYC* IRES structure Domain 1; only the base pairs in the hairpin spanning nt 110 to 136 (Fig. S3) are shifted over to allow the formation of pseudoknot helix α. Motif 9 partially overlaps Domain 2, where nt 284 to 299 of Domain 2 are refolded into two hairpins (Fig. S3). Structure models were compared vs. an alignment of 50 vertebrate *MYC* UTR sequences (Document S4). The alternative models for Doman 2/Motif 9 are roughly equally well supported by comparative data. Both are comprised of base pairs conserved across vertebrates and that show evidence of possible compensatory mutations: e.g. C279–G284 in Domain 2 vs. A307–U334 and G309–C332 in Motif 9 (Fig. S3). Neither model can be discarded based on these data. Nucleotides within Motif 9 were found to be highly reactive to chemicals in the previous *in vitro* analysis of the *MYC* UTR [6], thus their modeling as single stranded RNA. When overlaid on Motif 9, however, only 4 out of 21 modification sites were inconsistent with the `ScanFold-Fold` generated model (Fig. S3); additionally, sites of AMV reverse transcriptase pausing suggest that this region is structured.

Across the *MYC* mRNA, predicted structural metrics are most favorable in the windows that overlap Motif 17 in the 3’ UTR (Fig. 1). There are marked dips in the ΔG°, ED and z-score; all indicating importance of structure in this region. The `ScanFold-Fold` predicted base pairs in Motif 17 are also the best-conserved across the 15 vertebrate alignment. Previous work on post-transcriptional regulation of *MYC* found that inclusion of the short 3’ UTR sequence led to repression of luciferase expression [14] due to the inclusion of a miR-34 binding site. To determine if RNA structural features in the short 3’ UTR (beyond Motif 17) could be playing additional roles, the entire sequence was refolded while constraining Motif 17 base pairs. The resulting global short UTR model (Fig. 2) places the `ScanFold-Fold` predicted Motif 17 into a multibranch loop structure that includes a novel short hairpin. An additional hairpin is also predicted downstream of the multibranch loop. The short 3’ UTR model was analyzed vs. an alignment of 59 vertebrate *MYC* 3’ UTR sequences (Document S5). This found the highest levels of base pair conservation in the two long Motif 17 hairpins (92% conservation), while the remaining structures are not well-conserved (64% conservation of base pairing). When mutations occur in the highly-conserved Motif 17 they preserve base pairing: e.g. four compensatory (double point) mutations are found in each hairpin in addition to four and two consistent (single point) mutations, respectively (Fig. 2). To see if an orthogonal approach would confirm the 3’ UTR model structure or, perhaps, yield a better-conserved alternative model, the program RNAalifold [15] was used to evaluate the short 3’ UTR alignment without any base pairing constraints. The RNAalifold program considers both the folding energy and comparative sequence data (implicitly) in prediction; the resulting consensus model (Figure S5) predicts conserved structures that correspond to the two highly-conserved Motif 17 hairpins predicted by `ScanFold-Fold`.

**Figure 2.**
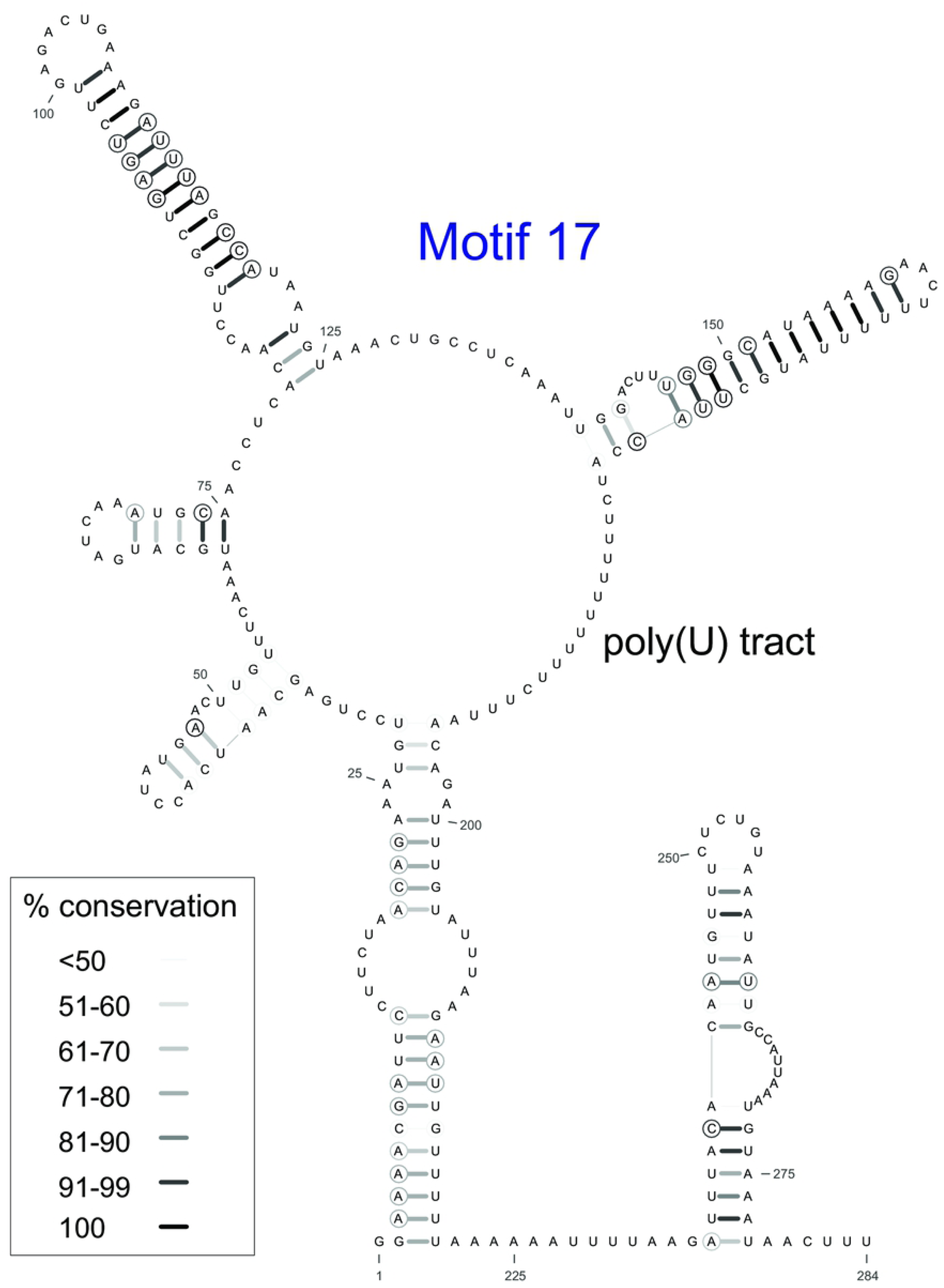
Short MYC 3’ UTR model. HP2–3 comprise Motif 17 of the `ScanFold-Fold` results, which were constrained in the calculation. Base pair conservation shading indicated in the key and data are taken from a comparison of 59 RefSeq mRNA vertebrate alignment (Document S5).

### Functional analyses of the MYC 3’ UTR

As the most significant motifs predicted in *MYC* occurred in the 3’ UTR, a known site of miRNA targeting, the locations of *MYC*-targeting miRNA binding sites were queried vs. predictions of structure. Of nine miRNAs with known interaction sites [8, 14, 16–20], *seven* occurred within Motif 17 (Fig. 3A and B). miR-34a/b/c, miR-449c and let-7a have overlapping seed binding sites in the unstructured region between the two highly-conserved Motif 17 hairpins (Fig. 3A and B). miR-145 binds downstream and partially overlaps the second hairpin. miR-148 has a seed binding site on the terminal stem-loop of the second hairpin. Interestingly, the two miRNAs that bind outside Motif 17 also do so in other `ScanFold-Fold` predicted structural motifs: miR-24 binds in the stem region of Motif 18 (Fig. 3C), while miR-185 binds toward the 5’ end of Motif 15 (Fig. 3D). In all cases, conserved RNA structures are predicted to partially occlude miRNA target binding, potentially modulating their effects.

**Figure 3.**
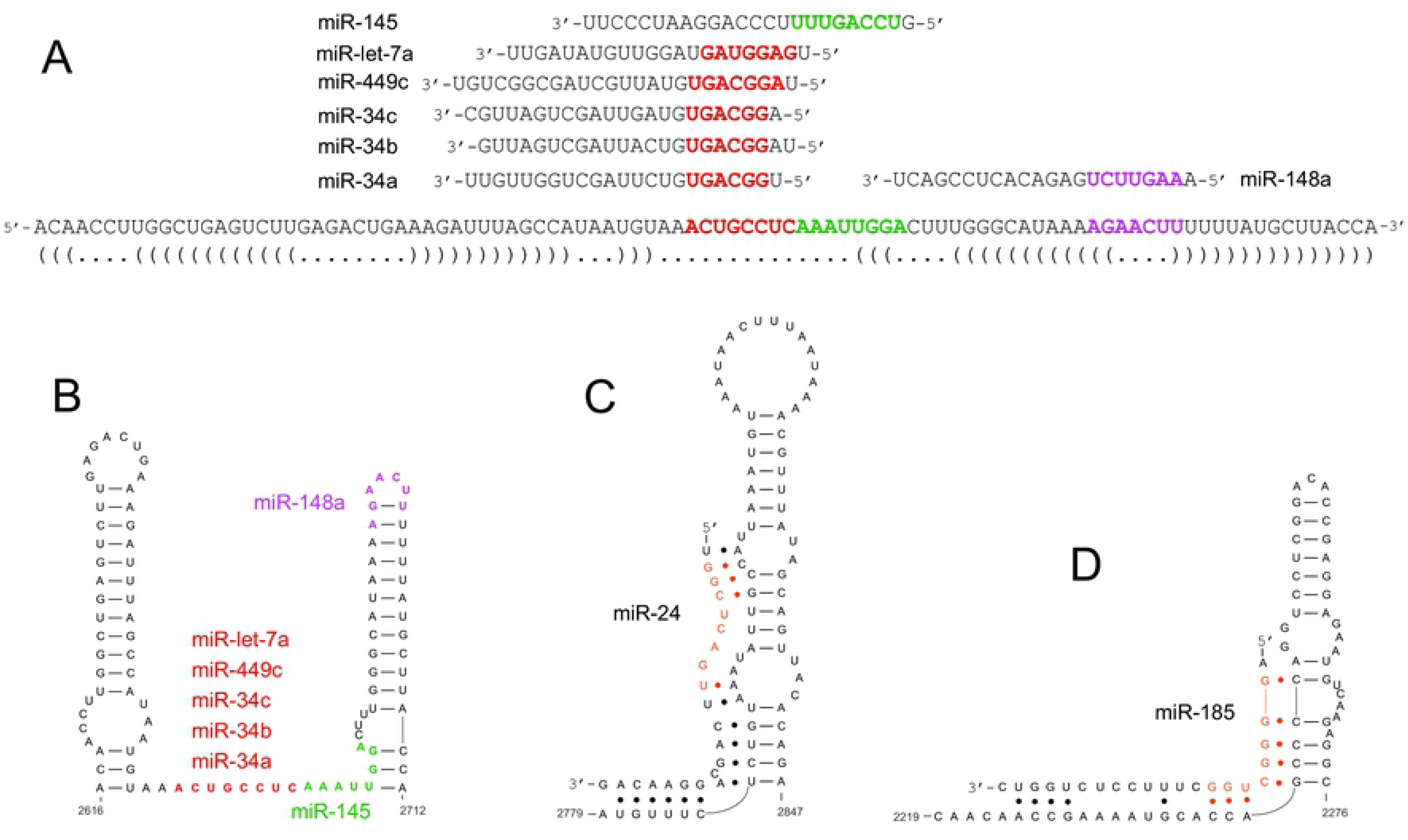
Annotations of miRNA binding sites on `ScanFold-Fold` predicted motifs. (A) Shows miRNA sequences above the “dot-bracket” structure of Motif 17 (matched brackets indicated base pairs). Seed sites and the complements on Motif 17 are colored. (B) Shows miRNA seed binding sites annotated on the 2D model of Motif 17. (C) Shows base-pairing between miR-24 and the 2D model of Motif 18. (C) Shows base-pairing between miR-24 and the 2D model of Motif 15.

Motif 17 was selected for additional experimental analysis due to it having the strongest `ScanFold` prediction metrics (Fig. 1), high level of structure conservation (Fig. 2), and the presence of multiple miRNA binding sites (Fig. 3A and B). To assess the potential gene regulatory roles of this motif, a luciferase reporter construct was generated incorporating Motif 17, along with 27 nt upstream and 11 nt downstream (including a poly(U) tract; Fig. 2). This sequence was inserted into the 3’ UTR of the <u>R</u>enilla luciferase (RL) expressing pIS2 vector (referred to as the pIS2-M17 [<u>M</u>otif <u>17</u>] vector; detailed in the Experimental Procedures). When assayed, the pIS2-M17 vector showed a significant decrease in both relative response ratio (RRR; Fig. 4)—a measure of luciferase activity—and translational <u>e</u>fficiency (TE) when compared to the unregulated pIS2 control: a ~24% and ~68% decrease in RRR and TE, respectively. This is consistent with previous analyses of the MYC 3’ UTR, where the entire short UTR isoform was incorporated into pLSV (an analogous Luciferase vector) and, using a similar analysis pipeline, was shown to lead to gene repression [14]. Similarly, ablation of the miR-34a-c seed (and also, seed regions for miRs 449c and let-7a) showed that miRNA targeting was responsible for the repressive effects of this region.

**Figure 4.**
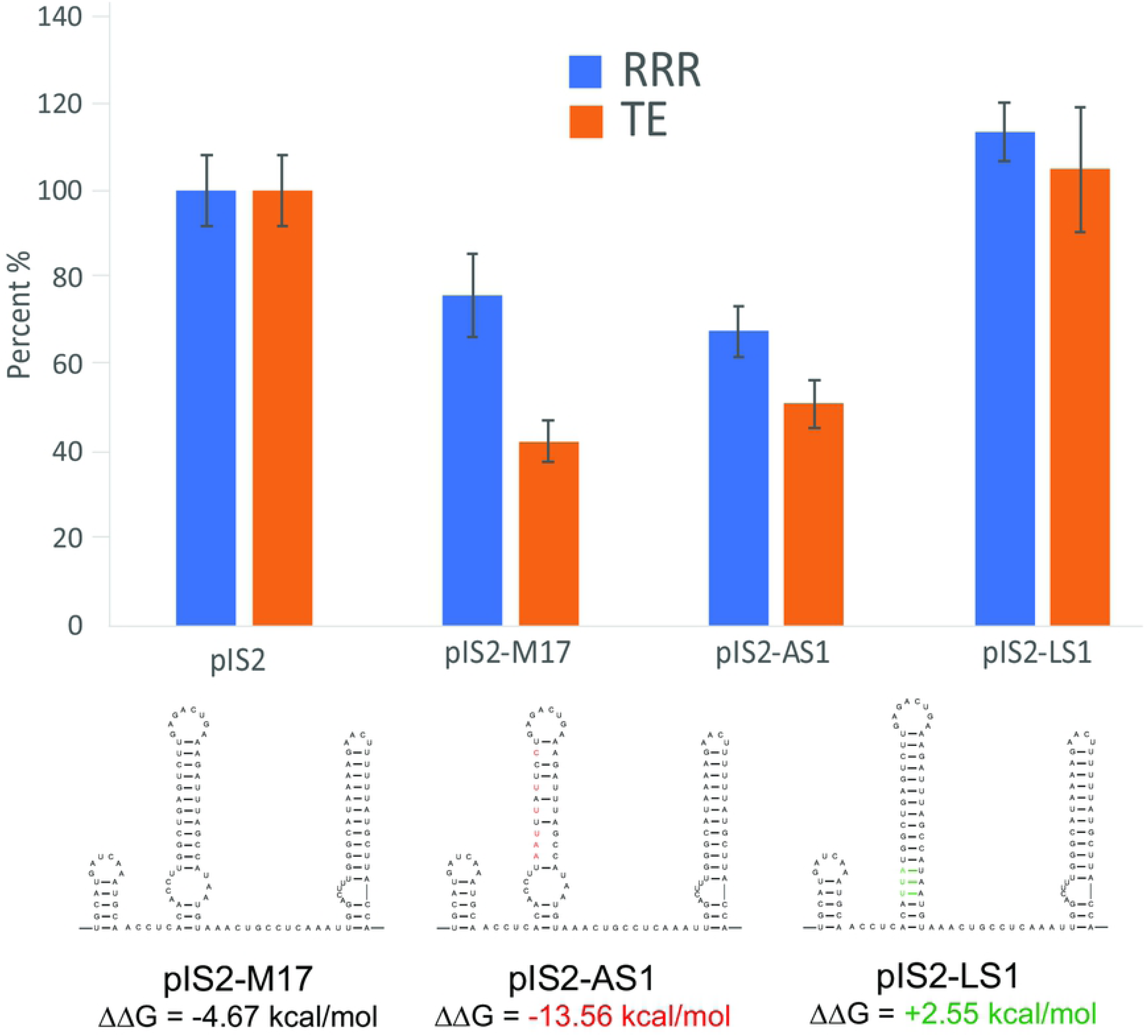
RRRs and TEs were calculated for each set of samples and normalized to the unregulated activity of pIS2. Error bars report the standard error. Experimental vectors pIS2-M17, pIS2-AS1, and pIS2-LS1 all display activity that differs from pIS2. pIS2-M17 and pIS2-AS1 both show decreased RRRs while pIS2-LS1, which was designed to have a more stable and less accessible 3’-UTR, shows an increase in RRR compared to pIS2. pIS2-MSC displays a large decrease in TE and pIS2-AS1, which is predicted to have a more accessible miRNA site, displays a TE which is slightly increased compared to pIS2-M17. The TE of pIS2-LS1 is markedly greater than pIS2-M17, possibly reflecting the decrease in miRNA target site accessibility. The ΔΔG values for pIS2-M17, pIS2-AS1, and pIS2-LS1 are also shown along with destabilizing pIS2-AS1 mutations (displayed in red) and stabilizing pIS2-LS1 mutations (displayed in green).

To determine if RNA structure present in Motif 17 influences miRNA binding/repression, two mutant constructs, pIS2-AS1 (<u>a</u>blate <u>s</u>tem <u>1</u>) and pIS2-LS1 (lock <u>s</u>tem <u>1</u>), were designed to increase or decrease miRNA site accessibility respectively (Fig. 4) according to the ΔΔG metric of Kurtesz et. al [21]. This metric accounts for both the energy needed to break native mRNA secondary structure and the energy gained by miRNA binding, and it was used to predict miRNA site accessibility for the WT and mutant constructs. The WT sequence, pIS2-M17, has a predicted ΔΔG of −4.67, whereas pIS2-AS1 and pIS2-LS1 have values of −13.56 (more accessible) and +2.55 (less accessible) respectively. When assayed, pIS2-AS1 shows (Fig. 4) a ~11% decrease in its RRR and an increase in TE of ~20% compared to pIS2-M17, however, these values are not significantly different from each other (Fig. 4). pIS2-LS1 showed a substantial increase in both RRR and TE (~68% and ~70% increase respectively) when compared to pIS2-M17.

## Discussion

The analyses performed in this report provide insights into the functions of RNA secondary structure in expression of *MYC*. The `ScanFold-Scan` results map out global features of RNA structure across the *MYC* mRNA. Interesting trends are observed moving across the sequence, where RNA structure thermodynamic stability decreases going 5’ to 3’, with marked “jumps” in instability observed at the UTR/coding-region junctions (Figs. S1 and S2). Likewise shifts toward more positive ED and z-score values were observed in junction-spanning windows: most dramatically at the 5’ junction, which includes both the CUG (non-canonical) and AUG (canonical) translation initiation sites. These results indicate a lack of stable structure here, reiterating previous observations that indicate inhibitory roles for thermodynamically stable RNA secondary structure at initiation sites [22]. We additionally find evidence that evolution may be specifically selecting for *MYC* initiation site sequences that are ordered to be less-stable than that predicted for sequences of similar composition (thus the positive z-scores); as well, the junction sequence is expected to have a volatile conformational ensemble, where no particular structure dominates (high ED).

The high and low respective thermodynamic stabilities of the 5’ and 3’ UTRs (Figs. S1 and S2) indicate differing roles for RNA folding in these regions. The highly-stable 5’ UTR would be expected to inhibit canonical translation by obstructing scanning ribosomes; thus, the presence of an IRES in the *MYC* mRNA. This can provide mechanisms for fine-tuning the post-transcriptional regulation of the *MYC* gene by allowing it to be translated in a cap-independent manner. The *MYC* IRES was shown to be active in some, but not all tissue types and the variability of activity is attributed to the presence, or lack of, trans-regulatory elements (e.g. RBPs; [4]). This demonstrates how cis-elements of the mRNA can interact with trans-regulatory elements to diversify (i.e. regulate) the cellular levels of a protein.

In contrast, the low stability of the 3’ UTR suggests a need for increased accessibility of the mRNA sequence: e.g. for intermolecular interactions with post-transcriptional regulatory factors such as miRNAs and regulatory proteins. Counterintuitively, the sites with the greatest evidence of having been ordered to fold into a specific structure are in the 3’ UTR (e.g. Motifs 17 and 18 in Fig. 1). Motif 17, for example, is the most well-conserved structured region in *MYC*—even more so than the IRES domain (Figs. 2 and S3)—and is supported via multiple compensatory and consistent base mutations. The highly favorable metrics and deep conservation of this motif throughout vertebrates indicated its biological importance, which was borne out by the analysis of miRNA binding sites (Fig. 3A and B) and Motif 17 function (Fig. 4). This motif acts as a hub for miRNA interactions and may organize the *MYC* miRNA target sites for interactions: e.g. the two highly-conserved hairpins may act as a “structure cassette” for maintaining the single-strandedness of the miR-34, −449c, and −let-7a seed binding regions, while modulating the accessibility of additional bases for miRNA pairing that can affect the outcome of miRNA targeting: e.g. translational repression.

The second most favorable motif (Motif 18; Fig. 1) contains a miR-24 interacting region (Fig. 3C). Notably, this interaction is “seedless” [17]—only three of the miR-24 seed nt are base paired to *MYC* (Fig. 3C). Most of the miR-24-interacting nt on *MYC* are predicted to be bound up in structure. Here, as in other interaction sites, RNA folding may be modulating accessibility. We observed that reducing the accessibility of miRNA binding in Motif 17 via pIS2-LS1 increased RRR and TE─potentially by reducing the amounts of miRNA-mediated gene repression. On the other hand, pIS2-M17 (the WT motif) and pIS2-AS1 (that ablates the first stem loop) had similar RRR and TE. Additional cellular factors may influence how these structures affect miRNA targeting: e.g. interactions with RNA binding proteins or post-transcriptional modifications can affect RNA folding. Thus, structural motifs and their additional interactions and alterations may be a way to fine tune the effects of miRNA regulation of *MYC*.

Conserved RNA structural motifs may also serve other functions in regulating *MYC* expression. Both Motif 17 and 18 occur in a particularly GC-poor and thermodynamically unstable region of the 3’ UTR (Fig. 1). This is due to long stretches of highly-conserved poly(U) (<u>u</u>ridine) tracts that occur between and around these structural motifs (Documents S3 and S5). Interestingly, poly(U) tracts have been found to stabilize mRNA sequences via structural interactions with the poly(A) (<u>a</u>denosine) tail [23]. Thus, an additional function of *MYC* 3’ UTR structural motifs may be to organize poly(U) tracts to facilitate interactions with the poly(A) tail. For example, in Figure 2 the poly(U) sequence is bulged out between the second Motif 17 hairpin and the basal stem of the predicted multibranch loop, which could allow the poly(A) tail to “wind” around this structure: e.g. in an analogous way to poly(U)-poly(A) interactions in viral and ncRNA stabilizing elements [24, 25]. Notably, our qPCR data does not indicate degradation of miRNA targeted RL transcripts. Instead, slight transcript accumulation is observed in experimental samples (which contain the poly(U) tract) compared to unregulated pIS2 samples (Table S1). Motif 17, in addition to regulating miRNA target accessibility, may be affecting transcript stability by modulating interactions of the poly(U) tract with the poly(A) tail of the mRNA. More investigation is needed to parse out these interactions.

Additional functional motifs are predicted beyond Motif 18 (Fig. 1), which may also be functionally significant. Interestingly, cancer-associated *MYC* translocations [26] can lead to UTR truncations that delete predicted motifs: potentially impacting function and contributing to *MYC* dysregulation. Likewise, seven predicted motifs fall within the *MYC* coding region, which may be functionally significant: e.g. by providing roadblocks for translation that can affect protein folding [27] or by affecting interactions with regulatory factors [28]. Notably, miR-185 targets a sequence that overlaps Motif 15 (Fig. 3D), which falls within the *MYC* coding region (Fig. 1). Awareness of the importance of miRNA targeting in coding regions is growing [29] and, presumably, additional *MYC* miRNA interactors remain to be discovered.

To conclude, this report provides valuable information on *MYC* mRNA secondary structure that has implications toward a better understanding of post-transcriptional gene regulation. In addition to providing global structural data, discreet local motifs with a high propensity for function are proposed, including a particularly interesting motif in the 3’ UTR that has been functionally validated. We showed Motif 17 possesses post-transcriptional regulatory function and, at the very least, this function is a result of structure regulated miRNA targeting. Our findings illustrate the utility of `ScanFold-Scan` and `ScanFold-Fold` in finding structured, regulatory motifs and highlight the important role of RNA secondary structure in the post-transcriptional gene regulation of *MYC* expression. This study provides a roadmap for further analyses of the structure/function relationships in the *MYC* mRNA and a framework for understanding other experimental results. For example, identified clinically significant sequence variants can be cross-referenced to these results to deduce their potential impact on RNA folding. Additionally, these results generate a large list of structural motifs that may be druggable targets [30, 31] for *MYC*, which is considered undruggable at the protein level [3].

## Materials and Methods

### In silico analyses

The *Homo sapiens MYC* RefSeq mRNA sequence was downloaded from the NCBI nt database (GenBank Accession: NM_002467.5). `ScanFold-Scan` was run using a single nt step size and window sizes of 70 (Document S1) and 120 nt (results with the longer window size were unchanged [data not shown], this 70 nt window was used in subsequent analyses). RNA structural metrics were calculated for windows using the RNAfold algorithm [32] using the Turner energy model [33, 34] at 37 °C. Z-score calculations were performed using the following equation (adapted from the approach of [13]):

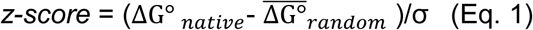

Here, ΔG°_*native*_ is the native sequence <u>m</u>inimum free <u>e</u>nergy (MFE) predicted by RNAfold. 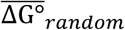 is the average MFE predicted for 100X mononucleotide randomized sequences. The standard deviation, σ, is calculated across all sequences. The other calculated data are: the P-value, which measures the fraction of random sequences that are more stable than native in the z-score calculation (this acts as a quality control measure for the z-score); the MFE ΔG°, which measures the thermodynamic stability of RNA secondary structure formation; the MFE base pairs that generate the MFE ΔG°, which are output in “dot-bracket” notation; the <u>e</u>nsemble <u>d</u>iversity (ED), which provides an estimate of the structural diversity in the RNA conformational ensemble (e.g. low ED indicates a single dominant conformation); the fraction of the (f)MFE in the ensemble, which estimates the contribution of the MFE conformation to the ensemble; the ensemble centroid structure, which is the conformation most similar to others in the ensemble; and the nt frequencies and GC percentages.

`ScanFold-Scan` prediction windows were next analyzed using the program `ScanFold-Fold` to deduce consensus motifs weighted by the z-score. The `ScanFold-Fold` method is detailed in [12]. Resulting output consisted of a list of all base pairing partners predicted for each nucleotide of the *MYC* mRNA (Document S6) and a list of the most favorable base pairing partners when weighting by z-score (Document S7). From the latter, base pairs which contributed to consistently negative z-scores (i.e. bps with average z-scores less −1 from Document S7) were used as constraints in an RNAfold prediction on the entire mRNA under the additional constraint of a maximum bp distance of 300 nt. Base pairs that extended `ScanFold-Fold` helixes were identified and used to generate the final motif models (Document S2). For visualizing the results of modeling, 2D rendering were generated using VARNA [35] and figures were produced with Adobe Illustrator.

For the analysis of conservation of `ScanFold-Fold` motifs across *MYC*, homologous mRNAs for 14 representative vertebrates were obtained from the NCBI RefSeq RNA database [36]. This database was also queried using BLAST [37] to deduce homologs for the “short” *MYC* 5’ and 3’ UTR sequences. Alignments for the mRNA (Document S3) and UTRs (Documents S4 and S5) were performed using MAFFT [38], implementing the MAFFT-E-INS-i and MAFFT-G-INS-i strategies, respectively [39].

A global model for the short 3’ UTR (defined/used in a previous study of miRNA targeting [14] was generated by constraining base pairs from Motif 17 and refolding the remaining sequence using RNAfold [32]. A consensus secondary structure for the short *MYC* 3’ UTR was predicted (Fig. S3) using RNAalifold [15] with the 3’ UTR alignment (Document S5) as input.

### Experimental analyses

#### Cell Culture

HeLa cells were incubated at 37°C and 5% CO_2_ and maintained in DMEM supplemented with 10% FBS, penicillin and streptomycin, and L-glutamine. Cells were passaged at 60-80% confluence and used between 3-40 passages.

#### Luciferase Vectors

For our experiments, two luciferase plasmid vector backbones were used. Both the transfection control vector, pIS0, which encoded firefly (FF) luciferase, and the experimental vector, pIS2, which encoded renilla (RL) luciferase, were gifts from David Bartel (Addgene plasmid # 12178; http://n2t.net/addgene:12178; RRID:Addgene_12178) and (Addgene plasmid # 12177; http://n2t.net/addgene:12177; RRID:Addgene_12177). The pcDNA3.1-miR34a vector was a gift from Heidi Schwarzenbach (Addgene plasmid # 78125; http://n2t.net/addgene:78125; RRID:Addgene_78125).

To test the post-transcriptional regulation of `ScanFold-Fold` predicted motifs, the Motif 17 sequence, along with 27 nt upstream and 11 nt downstream were incorporated into the 3’-UTR of pIS2 to generate pIS2-M17. Mutants that destabilize, pIS2-AS1, or stabilize, pIS2-LS1, the structure present in pIS2-M17 were generated. For pIS2-AS1, 6 mutations were incorporated that disrupt canonical base pairing in the first conserved hairpin. To generate pIS2-LS1, 3 mutations and 1 base deletion were introduced in the bulge on the upstream side of the first conserved hairpin. Mutations that destabilize or stabilize Motif 17 were predicted using the ΔΔG metric as a measure of miRNA site accessibility (Kertesz et al., 2007). 70 nt upstream and 70 nt downstream of the miRNA target site were included in our ΔΔG calculations.

The sequences for pIS2-M17, pIS2-AS1, and pIS2-LS1 were ordered as gBlocks from IDT and cloned using AgeI (5’) and Spe1 (3’) restriction sites (sequences in Supplemental Table S2).

Insertion of experimental sequences into the 3’-UTR of pIS2 required double restriction enzyme digest (using AgeI and SpeI from NEB) of both the gBlock and pIS2, following digestion, fragment and vector DNA were purified (Zymo DNA Clean and Concentrator kit), ligated (T4 Ligase from ThermoFischer), and transformed into DH5α-T1 competent cells using standard procedures. Carbenicillin selected colonies were cultured and plasmids were extracted (Qiaprep kit) and sequenced using an Applied Biosystems 3730xl DNA Analyzer.

#### Dual Luciferase Assay

Dual luciferase assays followed recommendations of an established method (Etten et al., 2013). In brief, the pIS0 vector (FF) is transfected at constant levels across all samples to serve as an internal control to which RL luciferase expression is normalized. All samples were run as biological triplicates. HeLa cells were counted using a hemocytometer and plated in a 24-well dish at a density of 50,000 cells per well. After 48 hours, cells were transfected using Lipofectamine 3000 (ThermoFischer) with 500 nanograms total plasmid DNA at a 1:1:8 ratio (pIS0:pIS2-based:pcDNA3.1-miR34a). Twenty-four hours later, cells were trypsinized, resuspended, and split into each of a 24-well plate for RNA analysis (1 ml) and a 96-well plate for the dual luciferase assay (0.2 ml). After another 24-hour incubation, cells in the 96-well dish were lysed, and luciferase activity was measured using Promega’s Dual Luciferase Reagent Assay kit on a Biotek Synergy 2 plate reader with a collection time of 10 seconds. <u>R</u>elative response ratios (RRR), the ratio of RL to FF relative light <u>u</u>nits (RLUs), were calculated for each sample and then normalized to the empty, unregulated pIS2 RRR. Cells from the 24-well plate were placed in TRIzol (ThermoFisher) and either stored at −80°C or immediately processed as below.

#### RNA Processing and qPCR Analysis

Cellular RNA was purified from samples in TRIzol using Zymo’s Direct-Zol RNA Miniprep kit. Purified RNA was then Dnase I treated (NEB) for 2 hours at 37°C and the resulting DNase-treated RNA was purified with Zymo’s RNA Clean and Concentrator kit. Reverse transcription was done using 1 microgram of purified RNA, random hexamers, and Superscript III (ThermoFisher).

Relative abundance of RL transcripts across samples were measured by qPCR, performed using PowerUp SYBR Green Master Mix on 1% cDNA input on an Applied Biosystems QuantStudio 3 instrument (ThermoFisher). Data were analyzed using the ΔΔCt method, where the relative abundance of RL transcripts in the samples were determined using the FF transcript as the reference gene. <u>T</u>ranslational <u>e</u>fficiencies (TE), a normalization metric (RRR/2^[−ΔΔCt_RL_]), were calculated for each sample. Primers used in qPCR were: RL FWD 5’-ggaattataatgcttatctacgtgc-3’; RL REV 5’-cttgcgaaaaatgaagaccttttac-3’; FF FWD 5’-ctcactgagactacatcagc-3’; and FF REV 5’-tccagatccacaaccttcgc-3’.

All data are available in the Supplemental Information. `ScanFold-Scan` and `ScanFold-Fold` are available for download from GitHub: https://github.com/moss-lab/ScanFold. RNAfold and RNAalifold are both bundled within the ViennaRNA package [32]: available at: https://www.tbi.univie.ac.at/RNA/.

## Acknowledgments

This research was supported by NIH/NIGMS grant R00GM112877 (WNM) and startup funds from the Iowa State University College of Agriculture and Life Sciences and the Roy J. Carver Charitable Trust (WNM) as well as R01-GM097455 (MDD).

## Supplemental Material

Supplemental material is available for this report.

